# Effects of different concentrations and combinations of antibiotics on the dynamics of intracellular transposition in *Escherichia coli*

**DOI:** 10.64898/2026.07.24.740473

**Authors:** Richard N. Goodman, Ellinor Shore, Michael S. M. Brouwer, Peter Nambala, Nicholas Feasey, Nina Langeland, Sabrina J. Moyo, Andrew Singer, Adam P. Roberts

**Author notes:** Corresponding author: Adam P. Roberts. Department of Tropical Disease Biology, Liverpool School of Tropical Medicine, Pembroke Place, Liverpool, L3 5QA, UK.

## Abstract

The use of antimicrobial compounds in humans, animals and in agriculture leads to environmental antimicrobial contamination through domestic, industrial and agricultural wastewater. Efforts have been made to perform environmental risk assessments based on the potential of these compounds to select for antimicrobial resistance (AMR) at certain concentrations in bacteria. This has resulted in predicted no effect concentrations (PNEC) which determine the minimum thresholds required to select for resistance. However, the effects of these compounds on intracellular transposition within bacterial isolates, a major driver of AMR, have not been previously assessed. Here, we assess the effect of differing sub-inhibitory concentrations of the third-generation cephalosporin, ceftriaxone, on the rate of intracellular transposition in combination with colistin and kanamycin. Two triple replicons systems (RS1 and RS2) were developed to assess this, each containing a chromosome, plasmid and entrapment vector. We show that sub-inhibitory concentrations of ceftriaxone exert hormetic effects on the intracellular transposition rate in RS1 and a steady linear increase in RS2. This defines a predicted no effect concentration for transposition (PNEC_T)_ for ceftriaxone as 320 ng/L in RS1 and 3200 ng/L in RS2. This provides a minimum threshold for the environmental impact of ceftriaxone on biological systems at the sub-cellular scale, which is applicable to industrial standards of waste management, where consideration of ecological impact is central.

## Introduction

Antimicrobial resistance (AMR), where microorganisms acquire the ability to resist the effect of antimicrobial compounds, is a major threat to human health. Recent studies have shown there were 4.95 million deaths associated with AMR in 2019 (Murray *et al*., 2022) and 4.71 million deaths associated with AMR in 2021 (Kariuki, 2024). A recent report by the World Health Organisation (WHO) highlighted an alarming increase in resistance rates in 104 countries, notably in Gram-negative bacterial pathogens such as *Acinetobacter* spp., *Escherichia coli*, *Klebsiella pneumoniae* and *Salmonella* spp., with approximately 1 in 6 laboratory-confirmed bacterial infections worldwide caused by resistant bacteria (World Health Organization, 2025). To address the issue of AMR, the One Health approach considers both antimicrobial usage (AMU) and AMR across interconnected sectors of humans, animals and the environment (McEwen and Collignon, 2018).

The use, misuse and overuse of antimicrobial compounds in healthcare (Boxall, Wilkinson and Bouzas-Monroy, 2022), veterinary care and personal care (Boxall *et al*., 2012) can lead to environmental contamination (Singer *et al*., 2016). The concentration of antimicrobial compounds found in the environment can be affected by the method of disposal and the amounts disposed. The chemical properties, degradability and dilution of the compounds also change the effect it has on the environment (Murray *et al*., 2024). Many environments contaminated with these antimicrobial compounds also contain microorganisms, where the contaminants can drive the selection of antimicrobial resistance, either through point mutations or horizontal gene transfer (Larsson and Flach, 2022; Cocker *et al*., 2025). Horizontal gene transfer can drive the spread of antimicrobial resistance between and within bacterial populations in environmental settings (Castañeda-Barba, Top and Stalder, 2024). Further to this, antibiotics can co-select for multiple AMR genes co-located on the same plasmid (Goswami *et al*., 2020). The assembly and mobilisation of these AMR gene arrays are frequently driven by genetic elements such as transposons and integrons (Castañeda-Barba, Top and Stalder, 2024). Therefore, it is important to assess the rate of intercellular and intracellular dissemination of mobile genetic elements (MGEs)when exposed to differing concentrations of antibiotic compounds.

In order to mitigate the effect of antimicrobial contamination in the environment there have been various studies to assess the effect of antimicrobial compounds on biological organisms. Efforts have been made to perform environmental risk assessments based on the potential of the compound to select for AMR at certain concentrations in bacteria (Murray *et al*., 2021) and fungi (Stanton *et al*., 2026). One method uses MIC data to determine predicted no effect concentrations (PNEC) (Bengtsson-Palme and Larsson, 2016) or predicted no effect concentrations for resistance (PNEC_R)_ (Murray *et al*., 2024) which determine the minimum thresholds required to select for resistance. These thresholds can then be used in pharmaceutical or industrial manufacturing waste management where they serve as a regulatory tool for industry to set concentrations which are permissible in the environment (World Health Organization (WHO), 2023). If the measured concentration exceeds the PNEC_R i_t could pose a risk to the environment by selecting for resistant bacteria over time. However, there are no current efforts to assess the effect of antimicrobials compounds on the transfer of MGEs and the AMR genes they often mobilise.

Entrapment vectors are molecular tools which can be used to detect intracellular transposition in the cell. They are designed so that insertion of an MGE affects the expression of a gene which can be phenotypically selected (Sołyga and Bartosik, 2004). Previously the pBACpAK entrapment vectors have been used to capture MGEs in various strains and isolates (Tansirichaiya *et al*., 2020, 2022, 2025; Goodman *et al*., 2023; Goodman, Tansirichaiya and Roberts, 2023). pBACpAK utilises a repressor-antibiotic resistance gene system with the *cI* gene, originally from λ phage, encoding the λ repressor protein, which inhibits transcription from the P_R p_romoter upstream of the *tetA* gene (Tansirichaiya *et al*., 2020). When an MGE inserts into the *cI* gene, it derepresses the *tetA* gene, resulting in phenotypic tetracycline resistance. Entrapment vectors provide a system for detecting the transfer rate of MGEs (i.e. transposition rate) and determining thresholds above which compounds exert an effect on MGE transfer.

In this study, we developed two replicon systems, these are *Escherichia coli* strains, each containing a fully sequenced and annotated chromosome, plasmid and entrapment vector. The only difference between the two replicon systems is the selectable marker in the entrapment vector, carrying either a colistin or kanamycin resistance gene. We used the pBACpAK entrapment vector derivatives pBACpAK-COL and pBACpAK-KAN (Goodman, Tansirichaiya and Roberts, 2023) in these replicons systems to determine the effect of differing concentrations (0ng/L, 32ng/L, 320ng/L and 3200ng/L) of the third-generation cephalosporin (3GC) ceftriaxone on the rate of transposition into pBACpAK entrapment vectors.

## Methods

### Strains, plasmids and growth conditions

Bacterial strains and plasmids used in this study are listed in Table 1. Primers used in this study are listed in Supplementary Table 1. All strains were grown in LB medium (Sigma-Aldrich, MO, USA) at 37°C, when mechanical shaking was required, we used 200 rotations per minute (rpm). Antibiotics (all from Sigma-Aldrich, MO, USA) were used in growth media at the following concentrations: chloramphenicol (CHL) 12.5 μg/ml, colisitin (COL) 1 μg/ml, kanamycin (KAN) 25 μg/ml, tetracycline (TET) 5 μg/ml. Ceftriaxone (CRO) sodium salt (Santa Cruz Biotechnology, TX, USA) was made to a 2 mg/ml stock solution then serially diluted to 10^−4^ in H_2O_. From the 10^−2^, 10^−3^ and 10^−4^ dilutions 16 µl ceftriaxone solution was added to 84 µl H_2O_ to make 3200 ng/ml, 320 ng/ml and 32 ng/ml solutions respectively. These were then diluted by 10^−3^ when added to LB broth to make final working concentrations of 3200 ng/L, 320 ng/L and 32 ng/L. For example, to make 10 ml LB broth supplemented with 32 ng/L, 10 µl of the 32 ng/ml stock solution was added to 990 µl of LB broth.

**Table 1:** Replicon systems used in this study. All replicon systems contained the *E. coli DH5a* chromosome. CHL; chloramphenicol COL; colistin KAN kanamycin.

| Replicon System number | Species | Replicons in system | Antibiotics in media | Number of replicons |
| --- | --- | --- | --- | --- |
| RS1 | *E. coli* | DH5α/pD25466/pBACpAK-COL | CHL (12.5 μg/ml)<br>COL (1 μg/ml) | Chromosome: 1<br>Plasmids: 2 |
| RS2 | | DH5α/pD25466/pBACpAK-KAN | CHL (12.5 μg/ml)<br>KAN (25 μg/ml) | Chromosome: 1<br>Plasmids: 2 |

### Replicon system development

We developed two *E. coli* strains with three fully sequenced and annotated replicons, which we called replicon systems (RS). Each replicon system contained the DH5α chromosome, the pD24566 plasmid and either pBACpAK-COL (replicon system 1) or pBACpAK-KAN (replicon system 2). The only difference between the two replicon systems is the selectable marker in the entrapment vector (either pBACpAK-COL or pBACpAK-KAN), carrying either a colistin (*mcr-1*) or kanamycin (*aph(3ʹ)-Ia*) resistance gene (Goodman, Tansirichaiya and Roberts, 2023).

The pD25466 plasmid was originally extracted from a *K. pneumoniae* strain (D25466) isolated in Malawi (Musicha *et al*., 2019) and AMR genes of note include a *catA1* gene and *bla*_TEM-1B,_ the latter being within a Tn*2* transposon. To develop the replicon systems, a filter mating experiment described previously (Brouwer *et al*., 2020) was carried out using D25466 as donor and sodium-azide resistant *E. coli* DH5α-Azi^R^ which resulted in the *E. coli* DH5α/pD25466 transconjugant. The pBACpAK derivates were used to transform DH5α*/*pD25466 in an electroporation experiment described previously (Goodman, Tansirichaiya and Roberts, 2023) resulting in the two replicon systems: DH5α/pD25466/pBACpAK-COL (RS1) and DH5α/pD25466/pBACpAK-KAN (RS2). The transposants which contain insertions into pBACpAK are named according to the replicon system (RS1), assay number (RS1.1), the concentration of ceftriaxone (CRO-320), and the transposant number (1 to 96), for example RS1.1-CRO-320-1.

### Exposing replicon systems to different concentrations of ceftriaxone

Replicon systems were retrieved from a −80°C freezer, streaked onto LB agar plates with the appropriate selection (see Table 1) and incubated at 37°C for 16-18 hrs. Colonies were picked and used to inoculate 10 ml of LB broth supplemented with the appropriate antibiotics (see Table 1) and varying concentration of ceftriaxone 0 ng/L, 32 ng/L, 320 ng/L and 3200 ng/L. This concentration range was chosen because the PNEC value for ceftriaxone is 32ng/L (Bengtsson-Palme and Larsson, 2016) and we wanted to measure the PNEC (32ng/L), up to two orders of magnitude above the PNEC (320 ng/L and 3200 ng/L), and a control at 0ng/L. These cultures were incubated at 37°C and 200 rpm for 16-18 hrs. The overnight culture was then plated onto T plates with tetracycline selection representing putative transposants i.e., LB agar containing CHL, COL and TET for RS1 and CHL, KAN and TET for RS2 as undiluted and in dilutions of 10^−1^, 10^−2^ and 10^−3^, then incubated at 37°C for 18 hrs. The overnight culture was also plated onto N plates without tetracycline selection representing total viable cells i.e., LB agar plates containing CHL and COL for RS1 and CHL and KAN for RS2 in dilutions of 10^−5^, 10^−6^ and 10^−7^ then incubated at 37°C for 18 hrs. Colonies were counted for both T and N plates to determine colony forming units per milliliter (CFU/ml) from which transposition frequencies were calculated. Up to 48 colonies were selected for molecular confirmation from each concentration of ceftriaxone (0 ng/L, 32 ng/L, 320 ng/L and 3200 ng/L) and patch plated onto LB agar plates containing CHL, COL and TET for RS1 and CHL, KAN and TET for RS2, then incubated at 37°C for 16-18 hrs. There were three biological replicates for each replicon system.

### Screening for MGE insertion into entrapment vectors with PCR

Up to 48 colonies were transferred from the patch plate into 200 µl of LB broth in 96-well plates containing the antibiotics specific to the replicon system (see Table 1) and 5 µg/ml of tetracycline. These cultures were incubated up to 64 hours at 37°C, as the number of cultures increased up to this time. Once growth was observed, DNA was extracted by a boilate method as follows; 100 µl of culture was transferred to 200 µl PCR strip tubes and centrifuged at 4,200 x g for 10 mins. The supernatant was removed, and the cell pellet was resuspended with 100 µl nuclease free H_2O_, this was heated at 95°C for 15 mins and 600 rpm. 10µl of this boilate DNA was used as the template in a PCR reaction. The 25 µl PCR reaction mix was made by adding 12.5µl 2× Redmix (Bioline, Meridian Biosciences, OH, USA), 1.25 µl fwd primer at 10µM, 1.25 µl reverse primer at 10µM and 10 µl boilate DNA (see Supplementary Table 1 for primer sequences). The PCR reaction used an annealing temperature of 57°C and an extension time of 3 min. The PCR products were visualised by gel electrophoresis either manually on a 1% (*w*/*v*) agarose gel or automatically using a Tapestation 4200 (Agilent, CA, USA) with the D5000 screen tape and reagents. If the amplicon, which included the *cI* repressor, was different in size from the wildtype amplicon size (1.35kb) or where no amplicon was visible, the isolates were taken forward for sequencing. This included whole amplicon sequencing of the PCR amplicons, whole plasmid sequencing and rolling circle amplicon (RCA) sequencing, described below. All sequencing was performed by Plasmidsaurus (San Francisco, California, USA) using Oxford Nanopore Technology (Oxford, UK).

### PCR Amplicon sequencing

The PCR products were purified using the Monarch Spin PCR & DNA Cleanup Kit (New England Biolabs, MA, USA) according to the manufacturer’s instructions for the DNA cleanup & concentration using 5x volume of Buffer BZ at the binding step. DNA was eluted in 15 µl Buffer EY. The purified PCR amplicon DNA was sent to Plasmidsaurus for sequencing.

### Whole Plasmid sequencing

Transposant colonies were taken from the patch plate and used to inoculate LB broth containing CHL, COL and TET for RS1 and CHL, KAN and TET for RS2 and incubated at 37°C and 200 rpm for 16-18 hrs. Plasmids were extracted from the cultures with either the Monarch Plasmid miniprep kit (New England Biolabs, MA, USA) or the Pureyield Midiprep system (Promega, WI, USA) according to the manufacturer’s instructions. For the Monarch Plasmid miniprep kit, 9.25 ml of bacterial cell culture was pelleted (10 mins at 2862 *x g*) and resuspended. To elute the DNA, 30 μl of elution buffer was used at 50°C. The plasmid DNA was sent to Plasmidsaurus for sequencing.

### Rolling circle amplicon (RCA) sequencing

Since pBACpAK is a single copy plasmid, the plasmid extract concentration (µg/ml) was occasionally too low for whole plasmid sequencing. To amplify the plasmid DNA, we used the phi29-XT rolling circle amplification (RCA) kit (New England Biolabs, MA, USA), where we ran 20 µl reactions containing 2 µl circular plasmid template DNA, 4 µl 5x phi29-XT reaction buffer, 2 µl deoxynucleotide (dNTP) solution mix, and 10 µl *cI*-*tetA* forward primers at 100 µM (see Supplementary Table 1). This was heated to 95°C for 3 mins to denature the DNA. After denaturation, 2 µl phi29-XT DNA polymerase was added to each sample. The samples were incubated in a thermocycler at 42°C for 2 hrs followed by 65°C to inactivate the DNA polymerase. The RCA products were sent to Plasmidsaurus for sequencing.

### Analysis of transposition frequencies

Putative transposants (PT) were defined as the CFU/ml of tetracycline-resistant colonies after exposure to varying concentrations of ceftriaxone. Transposants (T) were defined as the CFU/ml of tetracycline-resistant colonies which were subsequently confirmed by PCR and sequencing to have an MGE insertion into the *cI* - *tetA* region of the pBACpAK plasmid. Minimal transposants (MT) were defined as the CFU/ml of transposants, which represent only unique transposition events, corrected for clonal expansion. This was done to account for the Luria-Delbrück fluctuation effect (Luria and Delbrück, 1943) i.e., to prevent clonal expansion of a single transposition event early in the growth phase from artificially inflating the transposition frequency. Any identical Tn/IS insertions which inserted at the same base-pair coordinate within the same independent biological replicate (assay plate) were collapsed and treated as a single, independent transposition event. If we could not confirm the exact insertion site from sequence data the transposon was given an identical dummy value so that the filtering only retained one for each novel IS/Tn. Determining the proportion of transposants confirmed by sequencing and the minimal transposants corrected for clonal expansion, allowed us to calculate a percentage of colonies screened, these were applied as a correction factor to the putative transposant CFU/ml (PT) to give the transposant CFU/ml (T) and the minimal transposant CFU/ml (MT). Therefore, three transposition frequencies were calculated: the putative transposition frequency (PT/N), the transposition frequency (T/N), and the minimal transposition frequency (MT/N), where N was the total CFU/ml growing without exposure to tetracycline. Transposition frequencies were plotted as jitterplots and boxplots in R v 4.3.1 with ggplot2 v 3.5.2. The compare_means function of the ggpubr v 0.6.0 package was used for pairwise significance testing using the t-test, correcting for multiple comparisons with the Benjamini-Hochberg False Discovery Rate (FDR) method. Log_10 t_ransformation of the transposition frequency was carried out with the log_10 b_ase function, adding a pseudocount of 1e-8 to avoid the undefined logarithm of 0. An ANOVA was run on the log_10 t_ransformed frequencies using the aov base R function. A second-degree polynomial (quadratic) regression model was fitted to the transposition frequency data and the pseudo-log transformed concentration using geom_smooth from the ggplot2 v 3.5.2 package. Mean and standard deviation (SD) of the transposition frequencies was determined by ceftriaxone concentration using base functions. Log_2 f_old change (LFC) was calculated using the log_2 b_ase R function with the equation: log_2(_AT/C) where AT is the mean transposition frequency for the varying ceftriaxone concentrations (32, 320 and 3200 ng/L) and C is the mean transposition frequency for control (0ng/L ceftriaxone). The standard error of the mean (SEM) was calculated for each concentration of ceftriaxone using the equation SD/√NR, where SD was the standard deviation and NR was the number of replicates. The concentration dependent effect of ceftriaxone on CFU/ml was visualised by fitting a generalized linear model (GLM) with a Gaussian error distribution to the pseudo-log transformed concentrations using geom_smooth from the ggplot2 v 3.5.2 package. Kruskal-Wallis eta squared effect size was calculated using the kruskal_effsize function from the rstatix v 0.7.2 package. Graphs were assembled into multi-panelled figures using the patchwork v 1.3.1.9000 package. The R scripts used in the analysis of transposition frequencies can be found at https://rngoodman.github.io/CRO-concentrations-Tn-frequency.

### Bioinformatic analysis

The consensus whole plasmid sequence in fasta format was provided by Plasmidsaurus. Plasmid sequences were annotated by Plannotate (McGuffie and Barrick, 2021) with insertion sequence elements annotated using ISFinder (Siguier *et al*., 2006) and transposons using TnCentral (Ross *et al*., 2021) or MobileElementFinder (Johansson *et al*., 2020). Concentric circular plasmid plots were made by reorientating the plasmids to *oriV* and parsing the genbank to a csv file using custom python scripts (https://rngoodman.github.io/CRO-concentrations-Tn-frequency) then using the circlize v 0.4.16 R package to visualise the plots. The gggenes v0.5.1 R package was used to visualise linear plasmid plots.

Insertion site analysis was run on any long read sequenced plasmids, amplicons or RCA products which showed the entire insertion of the MGE. To mitigate for the Luria-Delbrück fluctuation effect (Luria and Delbrück, 1943), multiple identical IS insertions inserted at the same base-pair coordinate within the same independent biological replicate (assay plate) were collapsed and treated as a single, independent transposition event. The pBACpAK-COL and pBACpAK-KAN sequences were used as references, reorientated to start at the *repE* gene, and the genbank file was parsed to a csv file containing gene name, start and end using custom python scripts (https://rngoodman.github.io/CRO-concentrations-Tn-frequency). The plasmid sequences were also reorientated to *repE*, then aligned to the pBACpAK plasmid reference in Snapgene v3.3.4 to obtain the insertion sites in bp (from the start of *repE* gene). The “lollipop” plots were visualised in R, using dplyr to calculate heights, offsets, and directions based on 4 tiers, then ggplot2 was used with geom_arrow to visualise the genes as arrows and geom_point to visualise the insertion sites as lollipops. To enlarge genomic regions of interest containing multiple insertions coord_cartesian was used, defining the x limits. Subsequent statistical tests were run on the raw data and the data corrected for clonal expansion (as described previously). To determine if the insertion target sites were significantly dependent on the ceftriaxone concentration, a Fisher’s Exact Test with simulated p-values (based on 2,000 replicates) was utilised to evaluate categorical independence. To determine insertion site diversity across concentrations, the Shannon Diversity Index was calculated on the independent transposition events to measure both the richness and evenness of the insertion sites using the vegan v2.6.8 package in R. This was followed by sample-size-standardised rarefaction analysis to determine the effect of the number of samples sequenced on the number of unique insertion sites.

### Data, Materials, and Software Availability

All data and code to reproduce this analysis are available through GitHub: https://rngoodman.github.io/CRO-concentrations-Tn-frequency. The annotated plasmid sequences generated in this study have been deposited in the European Nucleotide Archive (ENA) at EMBL-EBI under the Study accession number PRJEB121544.

## Results

### The development of reproducible intracellular entrapment replicon systems

Two replicon systems (RS) were developed: DH5α/pD25466/pBACpAK-COL (RS1) and DH5α/pD25466/pBACpAK-KAN (RS2). Each contained a fully sequenced and annotated chromosome, plasmid and pBACpAK entrapment vector (Figure 1). MGEs of note include IS*1A*, IS*5*, IS*10R* and Tn*1000* in the DH5α chromosome, and IS*Kpn25* and Tn*2* (carrying *bla*_TEM-1B)_ in the pD25466 plasmid (originally isolated from a clinical strain in Malawi (Musicha *et al*., 2019)). These are closed systems where any insertion into the *cI* – *tetA* region on pBACpAK derivatives comes from one of the two other replicons present in the cell.

**Figure 1:**
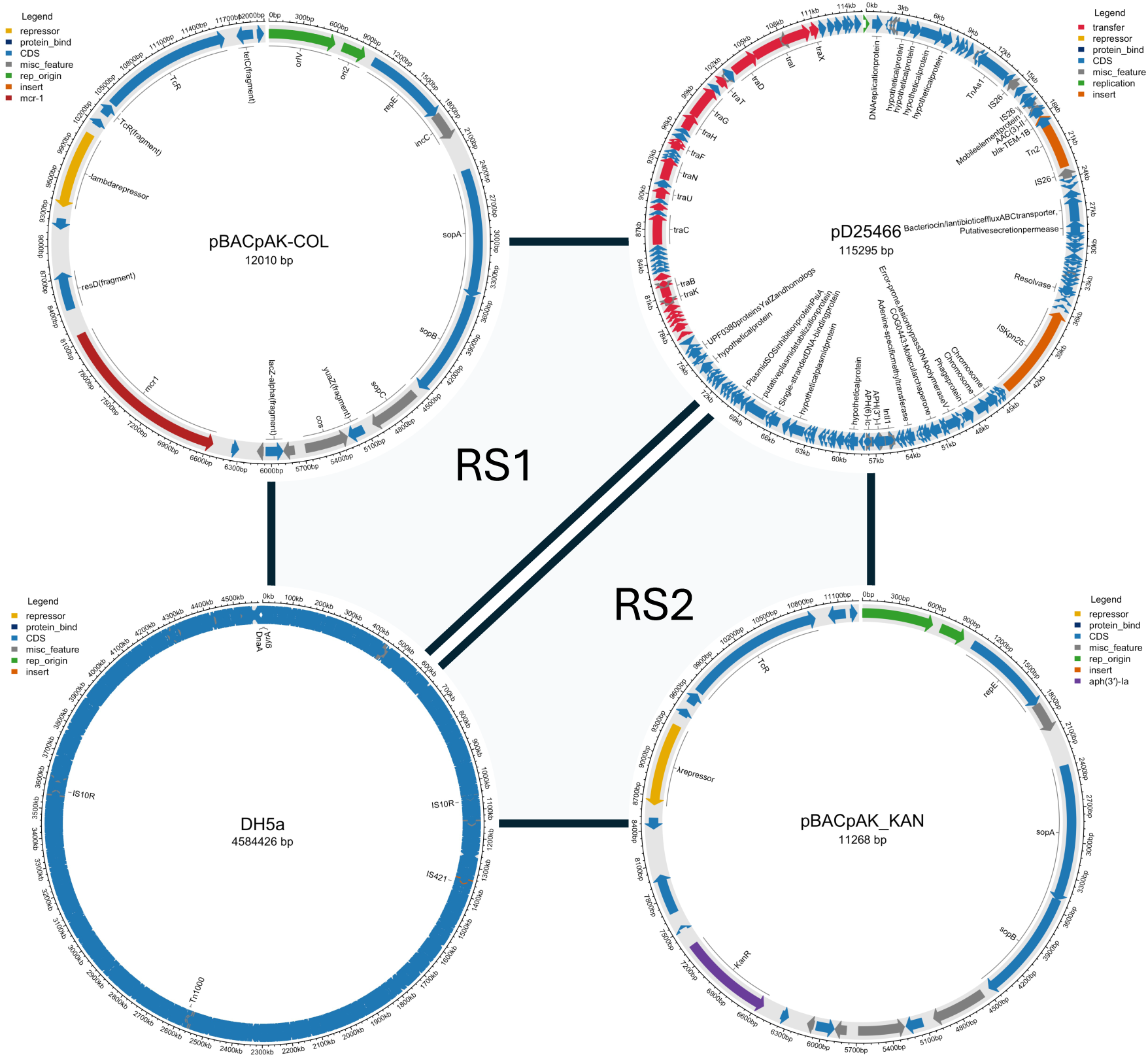
Annotated genome sequences of the replicon systems. Replicon system 1 (RS1) contains DH5α/pD25466/pBACpAK-COL, replicon system 2 (RS2) contains DH5α/pD25466/pBACpAK-KAN. The only difference is *mcr-1* (dark red) is present in RS1 and *aph(3’)-Ia* (purple) is present in RS2.

### Ceftriaxone exerts a dose-dependent inhibitory effect on *E. coli* growth

We determined CFU/ml at each ceftriaxone concentration to assess the impact of ceftriaxone on bacterial survival. Ceftriaxone was found to exert a dose-dependent inhibitory effect on *E. coli* growth for both RS1 and RS2 (see Figure 2). For RS1, at control conditions (0 ng/L of ceftriaxone) mean CFU/ml was 4.86 × 10^8^, the addition of 32 ng/L of ceftriaxone resulted in a modest reduction to 2.42 × 10^8^ CFU/ml. However, at a ceftriaxone concentration of 320 ng/L viable counts dropped to 8.7 × 10^7^ CFU/ml, representing a −2.48 log_2 f_old change (LFC) compared to control. At the highest concentration (3200 ng/L) the viable count plateaued at a similarly low level (7.7 × 10^7^ CFU/ml) indicating a - 2.66 LFC from control. For RS2, at control conditions (0 ng/L) the mean CFU/ml was 4.25 × 10^7^, reducing to 3.90 × 10^7^ at 32 ng/L (LFC: −0.125), then 2.19 × 10^7^ at 320 ng/L (LFC: −0.954), with the largest reduction to 1.47 × 10^7^ (LFC: −1.52) at the highest concentration of 3200 ng/L. Therefore overall, CFU/ml decreased with increasing ceftriaxone concentration.

**Figure 2:**
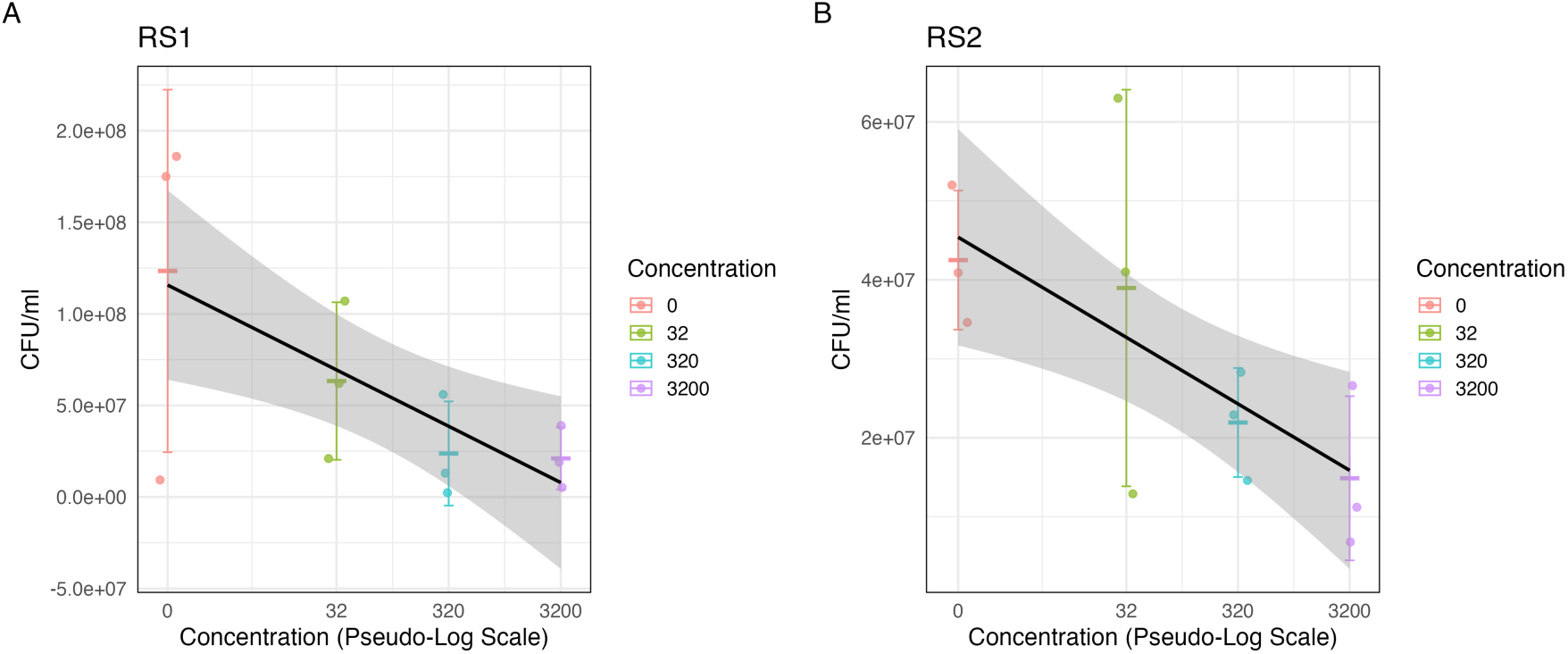
The effect of ceftriaxone on CFU/mL in replicon systems 1 and 2. Individual data points represent CFU/ml across the four concentrations of ceftriaxone (0ng/L, 32ng/L, 320ng/L, 3200ng/L) and three biological replicates. Horizontal crossbars indicate the mean CFU/ml for each group, with error bars representing one standard deviation above and below the mean. The solid black line represents a generalised linear model (GLM) fit to the data to illustrate the overall trend, with the shaded region indicating the 95% confidence interval. The x-axis is scaled using a pseudo-logarithmic transformation to visualise the 0ng/L concentration (using the pseudocount) and to equally space the concentrations (using the log transformation).

### Transposition occurs at low rates within our replicon systems

Colonies that grew on tetracycline supplemented agar plates, after incubation in differing ceftriaxone concentrations, were called putative transposants (PT). However, once confirmed by PCR and sequencing, we found the number of transposants (T) was much lower. Across all concentrations, an average of 30% of putative transposants were confirmed to be transposants by sequencing in RS1 (see Table 2) and 12% in RS2. The disparity came from isolates with point mutations or deletions within the *cI* repressor which also derepresses *tetA* and results in phenotypic resistance to tetracycline (Table 2). Therefore, transposition occurs at low rates within our replicon systems and the majority of phenotypic tetracycline resistance is accounted for by SNPs and INDELs in the *cI* repressor rather than insertion of MGEs.

**Table 2:** The percentage of transposants relative to putative transposants. Percentages calculated by dividing transposants (confirmed by PCR and sequencing) by total number of putative transposants (colonies growing on tetracycline plates). Mean calculated across 3 biological replicates.

| Replicon system | Concentration | Transposants confirmed by PCR/Sequencing (%) |  |
| --- | --- | --- | --- |
|  |  | Mean | SD |
| RS1 | 0 | 50 | 22.53 |
|  | 32 | 32.10 | 31.46 |
|  | 320 | 28.89 | 10.72 |
|  | 3200 | 9.80 | 16.98 |
| RS2 | 0 | 10.35 | 11.62 |
|  | 32 | 8.13 | 8.96 |
|  | 320 | 8.41 | 9.32 |
|  | 3200 | 22.07 | 22.61 |

### Ceftriaxone concentration affects the transposition frequency of mobile genetic elements in a biphasic, concentration-dependent manner in replicon system 1

We investigated the effect of ceftriaxone concentrations on the putative transposition frequency (PT/N) in RS1. We observed a non-linear, biphasic response to increasing ceftriaxone concentrations (Figure 3G). The largest increase was observed at a concentration of 320ng/L, the mean transposition frequency rose to 2.22 × 10^−6^ representing a 2.88 log_2 f_old increase relative to the control (include value). The ceftriaxone concentration of 320ng/L also exhibited the highest variance among biological replicates (Standard Error of the Mean [SEM] +/− 1.84 × 10^−6^; Supplementary Figure 1). In contrast, the highest concentration tested (3200 ng/L) resulted in a sharp decrease in transposition frequency to 5 × 10^−7^, which was only slightly higher than the baseline (0.731 log_f2_ old increase from baseline). An ANOVA found no significant difference across concentrations (*p* = 0.22) (Figure 3A).

**Figure 3:**
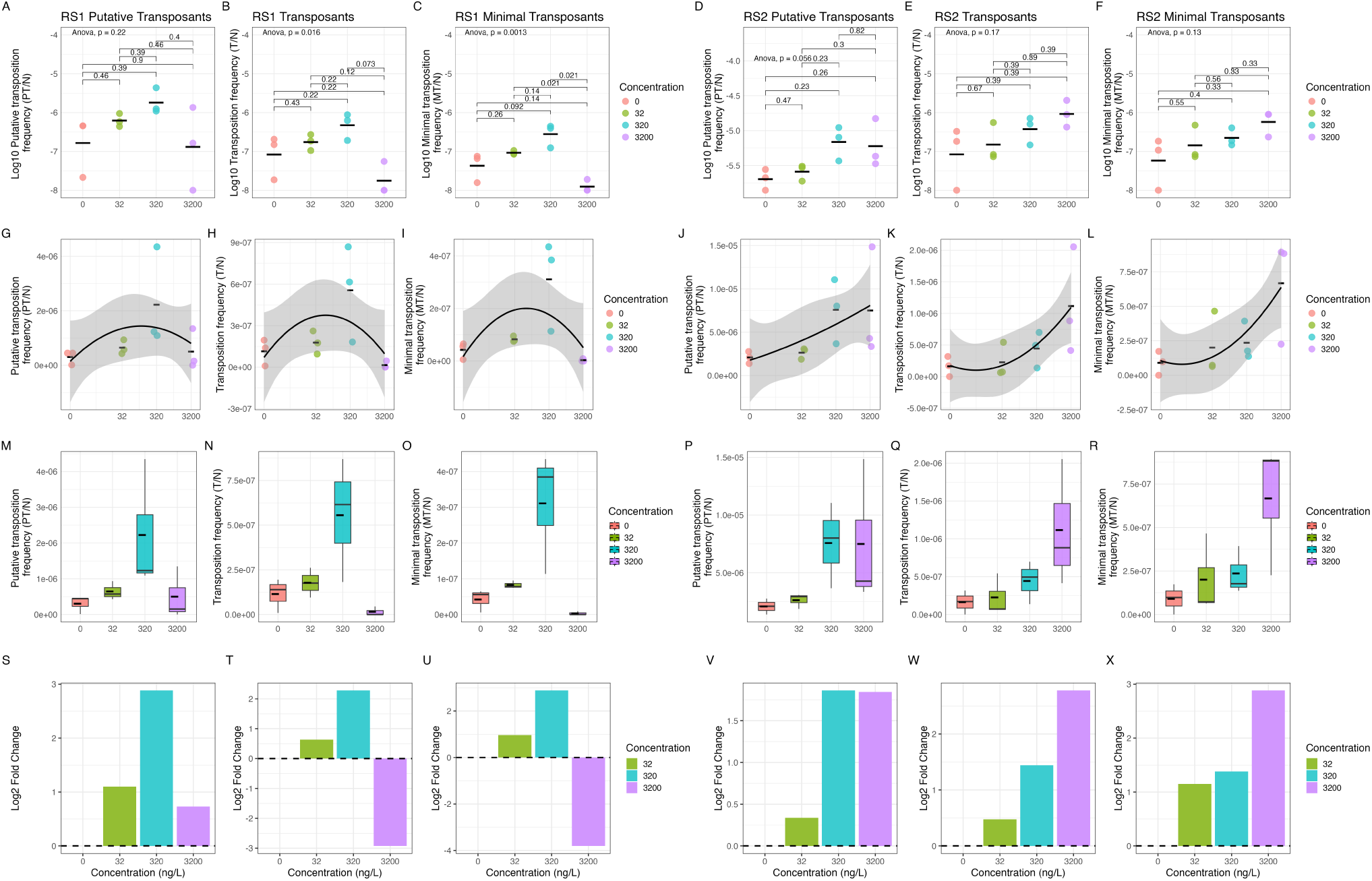
The effect of increasing ceftriaxone concentration on transposition frequency of MGEs within replicon systems (RS) 1 and 2. The x axis represents the ceftriaxone concentrations in ng/L. Putative transposants (PT) represent any colonies growing on tetracycline plates, transposants (T) represent insertions verified with PCR or sequencing, minimal transposants (MT) represents insertions verified with sequencing collapsed together based on their insertion site to ensure only unique insertion events are represented. A-F Log_10 t_ransformed transposition frequency of putative transposants (PT), transposants (T) and minimal transposants (MT) for replicon system 1 (RS1) and replicon system 2 (RS2). An ANOVA was carried out between all concentrations. Pairwise t-tests were carried out between each pair of concentrations (*n* = 6) using the Benjamini-Hochberg False Discovery Rate (FDR) method to correct for multiple comparisons and control the false discovery rate. G-L Linear regression showing effect of increasing concentrations of ceftriaxone on transposition frequency of PT, T and MT in RS1 and RS2. Horizontal crossbars denote the mean of each group. A second-degree polynomial linear regression model was fitted to the data (solid black line), with the shaded region indicating the 95% confidence interval. M-R Boxplots showing transposition frequency of PT, T and MT for RS1 and RS2. The median is represented by solid black line, the mean represented by dashed black line. S-X Log_2 f_old change relative to 0ng/L ceftriaxone (baseline) for 32, 320 and 3200ng/L of PT, T and MT for RS1 and RS2. The dashed line represents baseline.

We then screened the putative transposants for true insertions to determine the transposition frequency (T/N). At 0ng/L ceftriaxone the mean transposition (T) frequency was 1.14 × 10^−7^. We observed the same biphasic response to increasing ceftriaxone concentrations as seen with the putative transposition frequency (Figure 3H). The mean T frequency increases at 32ng/L (1.77 × 10^−7^; LFC: + 0.633) then is highest at 320 ng/L (5.55 × 10^−7^; LFC + 2.28) before a large decrease at 3200 ng/L (1.51 × 10^−8^; LFC: - 2.92) which was much lower than the baseline. Once again 320 ng/L had the highest variance across replicates (SEM: 2.01 × 10^−7^; see Supplementary Figure 1). An ANOVA found there was a significant difference between the transposition frequencies of different ceftriaxone concentrations in RS1 (*p* = 0.016) (Figure 3B).

Therefore, increasing ceftriaxone concentrations exerts a hormetic effect on the transposition rate (as shown by the “inverted-U” shape in Figure 3H). Concentrations of ceftriaxone from 0 to 320 ng/L cause an increase in transposition, 3200 ng/L is inhibitory, indicating that a threshold for cellular viability required for successful transposition events lies between 320 ng/L and 3200 ng/L.

### Ceftriaxone concentration affects the transposition frequency in a linear concentration-dependent manner in replicon system 2

We investigated the effect of ceftriaxone concentrations on the putative transposition frequency (PT/N) in RS2. Under control conditions (0 ng/L ceftriaxone) the mean spontaneous putative transposition frequency was 2.09 × 10^−6^. We observed a biphasic response with a peak frequency of 7.58 × 10^−6^ at 320 ng/L of ceftriaxone (+1.86 log_2 f_old change [LFC]) before dropping to 7.49 × 10^−6^ at the highest ceftriaxone concentration of 3200 ng/L (LFC: +1.84) (Figure 3J). The ANOVA was not significant (*p* = 0.056) (Figure 3D). The transposition frequency (T/N) exhibited a steady, linear increase in response to increasing ceftriaxone concentrations, up to the maximum tested dose of 3200 ng/L (Figure 3L). At baseline (0 ng/L) the mean transposition frequency was 1.63 × 10^−7^, this increased to 2.26 × 10^−7^ at 32 ng/L (+0.474 LFC), then 4.43 × 10^−7^ at 320 ng/L (+1.44 LFC) and 1.12 × 10^−6^ at 3200 ng/L (+2.78 LFC). Therefore overall, the transposition frequency of mobile genetic elements increases linearly with ceftriaxone concentration, however, there was no significant difference between the transposition frequencies of different ceftriaxone concentrations (*p* = 0.17; ANOVA) (Figure 3E). In RS2, 3200 ng/L has the highest variance across replicates (SEM: 4.88 × 10^−7^; Supplementary Figure 1), suggesting a heterogeneous population response to the ceftriaxone exposure.

This suggests increasing concentrations of ceftriaxone are inducing transposition but have not reached inhibitory concentrations at the maximum tested concentration of 3200 ng/L as in the case of RS1.

### Correcting transposition frequency for clonal expansion

With the sequence data we were able to determine the specific insertion sites for each MGE, this revealed that some transposition insertions were likely the result of clonal expansion following a single transposition event rather than each being a unique insertion event (also known as the Luria-Delbrück fluctuation effect) (Luria and Delbrück, 1943). To correct for this, we conservatively filtered to account for clonal expansion (see Methods) and applied the more conservative correction factor to the putative transposition frequency (PT/N) to give a minimal transposition frequency (MT/N). The minimal transposition frequecy (MT/N) had a larger log_2 f_old decrease than the transposition frequency (T/N) when compared to the putative transposants (Supplementary Figure 2). For RS1, the biphasic hormetic effect remained with increasing concentrations of ceftriaxone (Figure 3I), with an ANOVA finding a significant difference across different ceftriaxone concentrations (*p* = 0.0013) (Figure 3C). For RS2, the steady, linear increase also remained (Figure 3L) and there was still no significant difference across the concentrations (*p* = 0.13) (Figure 3F). Therefore, even when applying the most stringent correction factor using genomic sequencing data to correct for clonal expansion, the overall trends remain for RS1 and RS2.

### There was no significant difference in transposition frequency between replicon systems and the supplementation of colistin and kanamycin

We then determined the difference in all transposition frequencies between replicon system 1 and 2 (Supplementary Figure 3). Replicon system 1 (DH5a/pD25466/pBACpAK-COL) was supplemented with chloramphenicol (12.5µg/ml), colistin (1 µg/ml) and ceftriaxone (various), whereas replicon system 2 (DH5a/pD25466/pBACpAK-KAN) was supplemented with chloramphenicol (12.5µg/ml), kanamycin (25 µg/ml) and ceftriaxone (various). Therefore, the only difference between replicon systems is the resistance genes in pBACpAK, *mcr-1* in pBACpAK-COL of RS1 vs *aph(3ʹ)-Ia* in pBACpAK-KAN of RS2, or the supplementation of colistin (1 µg/ml) vs kanamycin (25 µg/ml). While there was a significant difference between the putative transposition frequency (PT/N) (*p* = 0.0069, pairwise t-test) of RS1 and RS2, there was no significant difference between the transposition frequency (T/N) (*p* = 0.15, pairwise t-test) and the minimal transposition frequency (MT/N) (*p* = 0.072) of RS1 and RS2 (Supplementary Figure 3) when considering all concentrations together.

### Defining a predicted no effect concentration for transposition (PNECT)

Whilst previous literature has presented a predicted no effect concentration for resistance (PNEC_R)_, no previous studies have focused on the effect of subinhibitory concentrations of compounds on intracellular transposition, therefore we propose a predicted no effect concentration for transposition (PNEC_T)_. In RS1, the lowest observed effect concentration (LOEC) for ceftriaxone was found to be 320 ng/L. Therefore, the PNEC_T f_or ceftriaxone would be 320 ng/L for RS1. Whereas in RS2, the lowest observed effect concentration (LOEC) for ceftriaxone was found to be 3200 ng/L, indicating an increase in the PNEC_T f_or RS2 to 3200 ng/L ceftriaxone.

### Genomic analysis of MGE insertion sites across the ceftriaxone concentrations

We analysed the genomes of the pBACpAK-COL and pBACpAK-KAN plasmids containing insertions across the different concentrations (Figure 4). IS*10R,* IS*1A,* IS*5,* Tn*1000* and IS*Kpn25* all inserted into the *cI* repressor at different locations, whereas Tn*2* inserted into both the *cI* repressor and *tetA* promoter (P_R)_, the inactivation of which also derepresses the *tetA* gene (Figure 4 and 5). IS*186B* was inserted within the *oriV* gene and was therefore an off-target insertion. In this case, phenotypic tetracycline resistance resulted from deletions in the *cI* repressor. Off target insertions, including those involving IS*186B,* were not included in the transposition frequency calculations.

**Figure 4:**
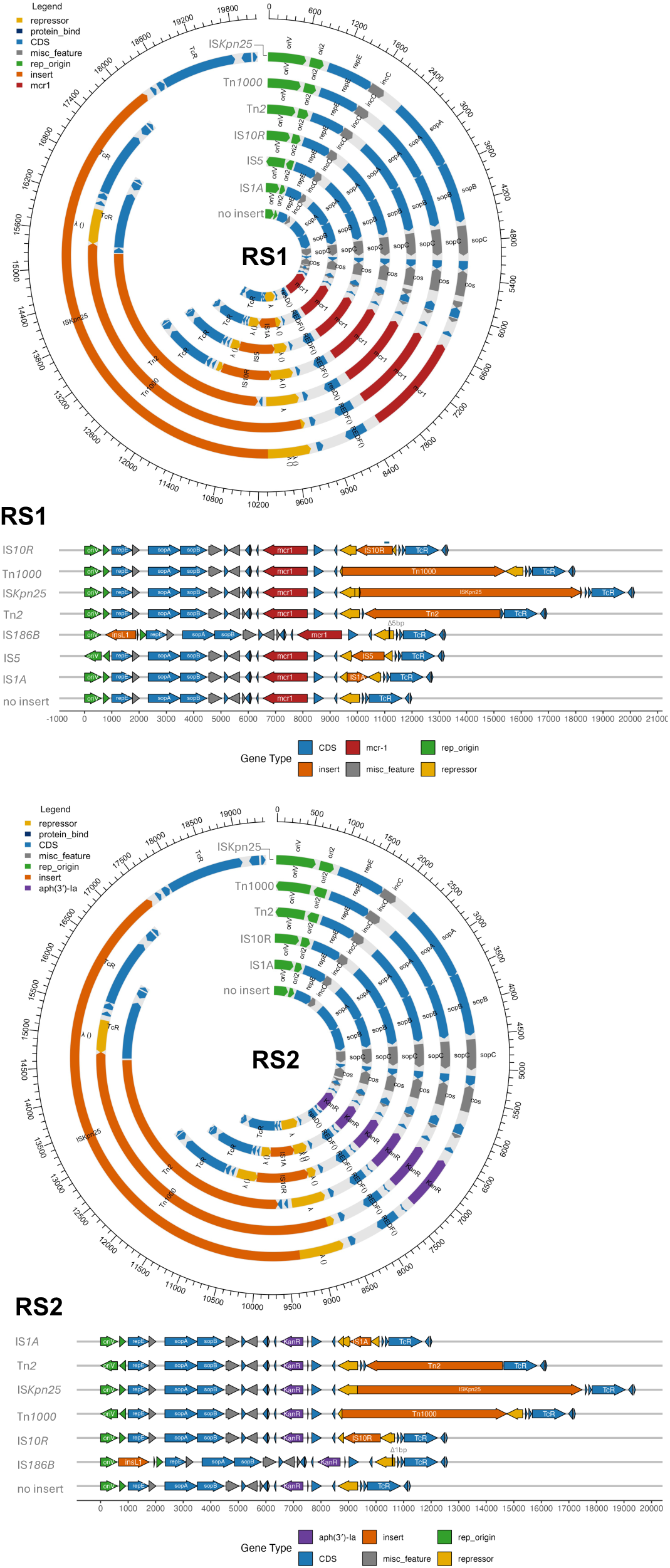
Genomic visualisation of IS and Tn insertions into pBACpAK-COL and pBACpAK-KAN. RS1 contained pBACpAK-COL, RS2 contained pBACpAK-KAN. All genomes were reorientated to start at the *oriV* gene. Deletions in the *cI* repressor are shown for IS*186B* which resulted in the phenotypic resistance to tetracycline in these transposants. These deletions are signified by Δ and the number of bp deleted when compared to reference (pBACpAK-COL or pBACpAK-KAN)

**Figure 5:**
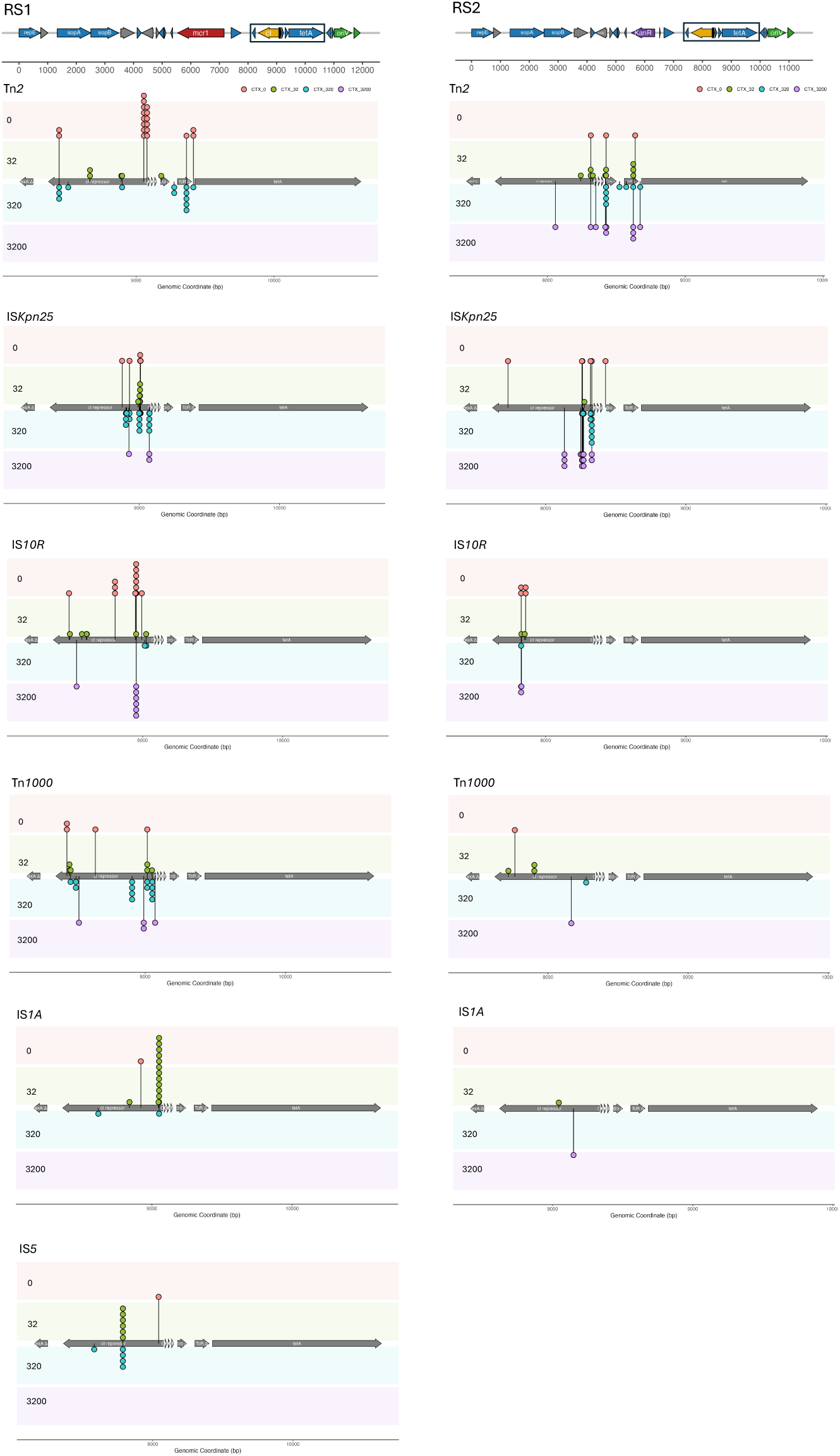
Comparison of insertion sites detected in pBACpAK-COL and pBACpAK-KAN. Both pBACpAK-COL (from RS1) and pBACpAK-KAN (from RS2) were reorientated to start at *repE*, therefore all basepairs are in relation to basepair 1 as the start codon of the *repE* gene. Both background panels and points are coloured according to ceftriaxone concentration in ng/L (0, 32, 320, 3200). Only fully sequenced plasmids, amplicons or RCA products containing a full MGE inserted into pBACpAK were included for this analysis. The uncorrected insertion sites are shown here, the insertion sites were collapsed to independent events for the diversity analysis (Shannon and rarefaction) and Fisher’s Exact Test.

We observed concentration dependent differences in insertion sites. The diversity of insertion sites for Tn*2*, IS*Kpn25* and Tn*1000* differ between ceftriaxone concentration in RS1 (Figure 5), where the highest diversity of insertion sites is at 320 ng/L for Tn*2* (H = 1.75 Shannon Diversity Index), IS*Kpn25* (H = 2.20) and Tn*1000* (H = 1.56) (Supplementary Figure 4). In the rarefaction analysis (Supplementary Figure 4), at the minimum common sample size, 320 ng/L does not have the highest richness for Tn*2*, IS*Kpn25* and Tn*1000*. This discrepancy between Shannon diversity (richness and eveness) and rarefaction (richness) shows that 320 ng/L seems to have a more evenly distributed insertion profile, indicating higher stochasticity at this concentration. Therefore, for Tn*2*, IS*Kpn25* and Tn*1000*, as the concentration increases in RS1, the insertion site diversity increases from 0 to 320 ng/L followed by a drop at 3200 ng/L (Supplementary Figure 4), which follows a similar trend to the hormetic curve of the transposition frequency. In RS2 the highest diversity of insertion sites is at 3200 ng/L for Tn*2* (Shannon Index, H = 1.97) and IS*Kpn25* (H = 2.08) (Supplementary Figure 4). Therefore, in RS2, the diversity of Tn*2* and IS*Kpn25* increases linearly with ceftriaxone concentration, again following a similar trend to the transposition frequency.

The insertion site analysis also revealed that clonal expansion was playing a part in transposition numbers, some insertion sites had several transposons inserted within the same assay and concentration (Figure 5). This is likely the Luria-Delbrück fluctuation effect (Luria and Delbrück, 1943), in which an insertion early in the broth culture’s growth phase has led to clonal expansion of the original event. Thus, when using the Fisher’s Exact Test, significant differences between concentrations were found for the insertion sites with the raw data; however, when the data was corrected for clonal expansion, these differences were no longer significant. There were significant differences in insertion site between the different MGEs in both RS1 (Fisher’s Exact Test, *p* = 0.0004998) and RS2 (*p* = 0.0004998).

## Discussion

In this study, we used an entrapment vector system to show that, in RS1, increasing concentrations of ceftriaxone exert significant hormetic effect on the transposition frequency. Whilst in RS2, there was a steady linear increase in transposition frequency as ceftriaxone concentration increased. The lowest observed effect concentration (LOEC) for ceftriaxone was 320 ng/L for RS1 and 3200 ng/L for RS2. This provides a minimum threshold for the environmental impact of ceftriaxone on biological systems measured at the sub-cellular scale. This threshold is applicable to industrial standards of waste management, where consideration of ecological impact is central. Whilst many studies have determined how differing concentrations of compounds have affected conjugation (Ding *et al*., 2022; Xiao *et al*., 2022; Alav *et al*., 2024; Zhou *et al*., 2024) this is the first to show PNEC associated subinhibitory dose response curves for transposition frequencies using entrapment vectors and also provide a predicted no effect concentration for transposition (PNEC_T)_.

In RS1, containing pBACpAK-COL and selected for with colistin, the transposition frequency (T/N) and the minimal transposition frequency (MT/N) were shown to exhibit a biphasic response to increasing ceftriaxone concentrations as shown in the hormetic (inverted-U) profile (Figure 3H and 3I) as described in previous literature (Davies, Spiegelman and Yim, 2006). The ceftriaxone concentration of 320 ng/L exhibited the highest transposition frequency, representing the highest mobile genetic element transposition. This is likely due to the increasing levels of stress on the bacterial cells, resulting in the activation of the SOS response (Baharoglu and Mazel, 2014), which triggers homologous recombination required for the DNA repair of double-strand breaks (Gillings, 2013; Chow, Waldron and Gillings, 2015), whilst remaining low enough for cell survival. This SOS response has been shown to be activated by beta-lactam antibiotics in *E. coli* (Miller et al., 2004) and *Staphylococcus aureus* (Maiques et al., 2006), thereby transiently halting bacterial cell division to avoid lethal antibiotic exposure. This stress response, caused by exposure to antibiotics, can lead to an increase in transposition rate (Vandecraen *et al*., 2017; Wagner *et al*., 2023), which has been coined “transposition bursts”(Wu, Aandahl and Tanaka, 2015). For example, subinhibitory concentrations of ciprofloxacin (0.2 µg/ml) and vancomycin (0.2 µg/ml) increased the transposition frequency of IS*256* in *S. aureus* (Nagel *et al*., 2011). Similarly in *E. coli*, sub-inhibitory concentrations of ceftazidime (0.5 µg/ml), cefotaxime (8 µg/ml), and piperacillin (64 µg/ml) were shown to enhance the transposition of IS*Ecp1B* associated with certain *bla*_CTX-M g_enes (from *Kluyvera* spp.) by 100-fold, 10-fold and 4-fold respectively (Lartigue *et al*., 2006). Here in this study, we have shown that increasing subinhibitory concentrations of ceftriaxone increases transposition frequency in *E. coli*, likely due to stress-induced transposition bursts.

The higher ceftriaxone concentration of 3200 ng/L indicates toxicity in RS1. There could be several reasons for this, such as saturation of DNA repair or overproduction of reactive oxygen species (ROS). Beta-lactam antibiotics have been shown to induce the SOS response in *E. coli* (Miller *et al*., 2004; Laureti, Matic and Gutierrez, 2013), which regulates the production of the *dinB* gene (Baharoglu and Mazel, 2014), which is cytotoxic to *E. coli* when overproduced (Foti *et al*., 2012). Therefore the higher concentration of ceftriaxone could force DNA polymerase, such as DinB (DNA Polymerase IV), to increase activity, targeting areas outside of those that are damaged, which could lead to lethal mutagenesis (Negishi, Loakes and Schaaper, 2002). It has also been shown that antibiotics induce redox stress, including the production of ROS, at higher concentrations of ceftriaxone the ROS could be overproduced and overwhelm the cells antioxidant defences (Dwyer *et al*., 2014). This cytotoxicity is likely why in RS1 the transposition drops well below baseline at 3200 ng/L. It is also interesting that the highest variance in insertion location amongst biological repeats was seen at 320 ng/L (shown by the highest SEM in Supplementary Figure 1). This could show that this concentration is near a tipping point where the fitness cost is more evenly balanced between the cost and benefits of transposition, or that the benefit from creating a fitness advantage from a positive transposition is more likely to be offset by cost of deleterious transposition events. This stochasticity is common in stress responses (Shimoni *et al*., 2009). Since there is uncertainty as to the time that bacterial cells spend being exposed to environmental stressors such as antibiotics, stochastic fluctuations in the stress-response of individual cells can dictate whether a population survives or dies (Sampaio *et al*., 2022). That is, if individual cells follow different evolutionary paths leading to heterogenous phenotypes, they can increase the chances that one of the cells will survive and reproduce once the environmental stressor has been removed from the environment. It is also worth considering the effect increasing beta-lactam concentrations, and any subsequent SOS responses, have on SNPs and INDELs. The beta-lactam induced activation of the SOS response (Baharoglu and Mazel, 2014) also leads to an increase in error prone polymerases, resulting in more SNPs and INDELs (Ahmad *et al*., 2025). Beta-lactam antibiotics can activate the RecA protein which represses *lexA*, allowing several genes associated with the SOS response to be expressed, including the error-prone polymerase DNA Polymerase IV (DinB) (Maiques *et al*., 2006; Galhardo *et al*., 2009). When exposed to sub-inhibitory concentrations of beta-lactams, the resulting polymerase-driven increase in SNPs and INDELs accelerates the exploration of the fitness landscape (Gutierrez *et al*., 2013; Laureti, Matic and Gutierrez, 2013). Thus, to observe a transposition event in a high-stress environment, a single cell must undergo a successful transposition event and simultaneously avoid a lethal SNP everywhere else in its genome. This assay assessed the transposition rate of the survivors, i.e., viable cells growing on tetracycline. This may have missed the cells with the most intense SOS response which had both higher transposition rates and a higher mutational load. This may have unintentionally introduced a selection bias where we observe a drop in transposition at higher concentrations of ceftriaxone (such as 3200 ng/L in RS1) because the cells that had both high SNPs and transposition didn’t survive.

In replicon system 2, containing pBACpAK-KAN and selected for with kanamycin, there is a gradual linear increase in transposition frequency with increasing ceftriaxone concentration to 3200 ng/L (Figure 3H), however the change is not statistically significant. In RS2 the concentration of 3200 ng/L has the highest variance across replicates (Supplementary Figure 1), which showed a heterogeneous population response similar to 320 ng/L in RS1. This suggests that increasing concentrations of ceftriaxone are inducing transposition, likely via stress-response pathways as mentioned previously (Baharoglu and Mazel, 2014), but have not reached inhibitory concentrations at the maximum tested concentration of 3200 ng/L as in the case of RS1. It could be that higher concentrations of ceftriaxone in RS2 would follow a similar curve to RS1 if the concentrations of ceftriaxone increased beyond 3200 ng/L. The reason toxicity may impact the bacterial populations at lower concentrations in RS1 compared to RS2 is likely due to the mechanism of action of the different antibiotics used for selection in these two systems: colistin and kanamycin respectively. Colistin is a polymyxin which acts on the cell membrane, binding to lipopolysaccharides (LPS) which disrupts membrane integrity (Moubareck, 2020), whereas kanamycin is an aminoglycoside antibiotic which binds to the 30S ribosomal subunit which disrupts polypeptide synthesis (Darby *et al*., 2022). While both antibiotics are bactericidal, at sub-inhibitory concentrations, their differing mechanisms may be the reason there are different observed effects from the increasing concentrations of ceftriaxone.

Whilst both our replicon systems contain resistance genes, *mcr-1* and *aph-3’-Ia*, the presence of antibiotics is still likely to cause physiological stress and potentially higher fitness costs. Colistin is more physically disrupting for the bacterial cell and therefore may make the replicon system more sensitive to increasing concentrations of ceftriaxone, so that 3200 ng/L is enough to dramatically reduce intracellular transposition in RS1, but not in RS2. The steadier increase in transposition frequency seen in RS2 (Figure 3K) could also be due to the mechanism of kanamycin, as small amounts which evade resistance, could be blocking the translation of transposase genes and masking transposition events. While this difference is an artefact of the experimental conditions of the replicon systems used in the entrapment vector assays, it shows that exposure to different antibiotics at both fixed and variable concentrations can all have differing effects on transposition rate within the cell. This is relevant to wastewater environments where antimicrobial compounds will dissolve and dilute at differing concentrations providing undulating niches for bacteria (Stanton *et al*., 2022). Many environmental conditions contain nutrient and chemical gradients such as sewage systems and rivers (Cocker *et al*., 2025). The fact that ceftriaxone concentrations as small as 320 ng/L can increase transposition frequency shows that even very small levels of contamination, from inappropriate disposal of antibiotics, can fundamentally affect biological systems.

Alongside MGE movement we also observed the transposition of the beta-lactamase gene *bla*_TEM-1B,_ as part of Tn*2*, which confers resistance to penicillin antibiotics. Insertion sequence mediated amplification of *bla*_TEM-1B h_as previously been shown to confer resistance to piperacillin/tazobactam through inhibitor saturation (Hubbard *et al*., 2020). Since Tn*2* moves via a “copy and paste” mechanism (replicative transposition) (Nicolas *et al*., 2015), the increase in Tn*2* movement at 320 ng/L of ceftriaxone (Figure 3), could drive an amplification of *bla*_TEM-1B w_hich in turn could drive resistance to beta-lactamase inhibitors such as tazobactam. Further to this, in some cases, the target site of Tn*2* was the P_R p_romoter, upstream of the tetracycline resistance gene, *tetA,* which derepressed expression of the gene by preventing the λ repressor from acting on the promoter. There have been other examples of IS elements inactivating resistance genes by inserting into promoters such as the silencing of *catA1* in *E. coli* through IS*5* inserting into the upstream promoter (Graf *et al*., 2024) and the silencing of *vanA* in *Enterococcus* spp. through IS*L3*-family element insertion into the upstream promoter (Sivertsen *et al*., 2016). Therefore, environmental pollution of antibiotic compounds could intentionally be driving antimicrobial resistance through the amplification of resistance genes or the inactivation of promoters driven by concentration-sensitive MGEs such as Tn*2*.

This study also shows how entrapment vectors and the triple replicon systems used here provide an adaptable platform for investigating the impact of various environmental stressors on mobile element and associated AMR dynamics. The *cI* gene, which encodes for the λ repressor, is originally from bacteriophage λ where it’s natural function is to maintain the lysogenic state by repressing genes involved in the lytic cycle (Atsumi and Little, 2006). The λ repressor and its homologues have been shown to be inactivated during stress response leading to the activation of integrative conjugative elements (ICEs) (Bellanger *et al*., 2007). Therefore, it is likely that many transposons and IS elements have evolved alongside bacteriophage λ with target sites in the *cI* gene.

This study has limitations. Up to 48 putative transposants were picked as single colonies from the tetracycline plates, introducing selection bias, as we may have missed colonies containing transposants that could have altered the transposition frequencies.

For the insertion site analysis, only sequenced plasmids, amplicons or rolling circle amplification products were used which did not represent all insertions and therefore may have biased the results. Targets outside of the *cI* – *tetA* region of pBACpAK (i.e. offsite targets) were not considered in the transposition frequency calculations (such as the insertion of IS*186B* into *oriV*), which may have led to under reporting of transposition frequency. To determine off-site targets, whole genome sequencing would have been required for every transposant which would not have been feasible.

To conclude, this study shows that sub-inhibitory concentrations of ceftriaxone exert a statistically significant hormetic effect on transposition frequency in RS1 and a steady linear increase in transposition frequency in RS2. We also defined a new PNEC, the PNEC_T,_ which is 320 ng/L ceftriaxone in RS1. The reproducible replicon systems we have developed provide an adaptable platform for systematically investigating the impact of various environmental stressors on mobile element and associated AMR dynamics. Furthermore, this study also provides a minimum threshold for the environmental impact of ceftriaxone, together with colistin and kanamycin, on intracellular MGE transfer within biological systems, which is applicable to industrial standards. This shows that even low, sub-inhibitory concentrations of an antibiotic in the environment can accelerate the evolution of AMR.

## Acknowledgements

This project (STRESST) received funding from the Medical Research Council (MRC), Biotechnology and Biological Sciences Research Council (BBSRC) and Natural Environmental Research Council (NERC) which are all Councils of UK Research and Innovation (Grant no. MR/W030578/1) under the umbrella of the JPIAMR - Joint Programming Initiative on Antimicrobial Resistance. A.P.R. acknowledges additional funding from the AMR Cross-Council Initiative through grants from the MRC and the National Institute for Health Research (grants no. MR/S004793/1 and NIHR200632).

## Supplementary Material

**Supplementary Table 1:**
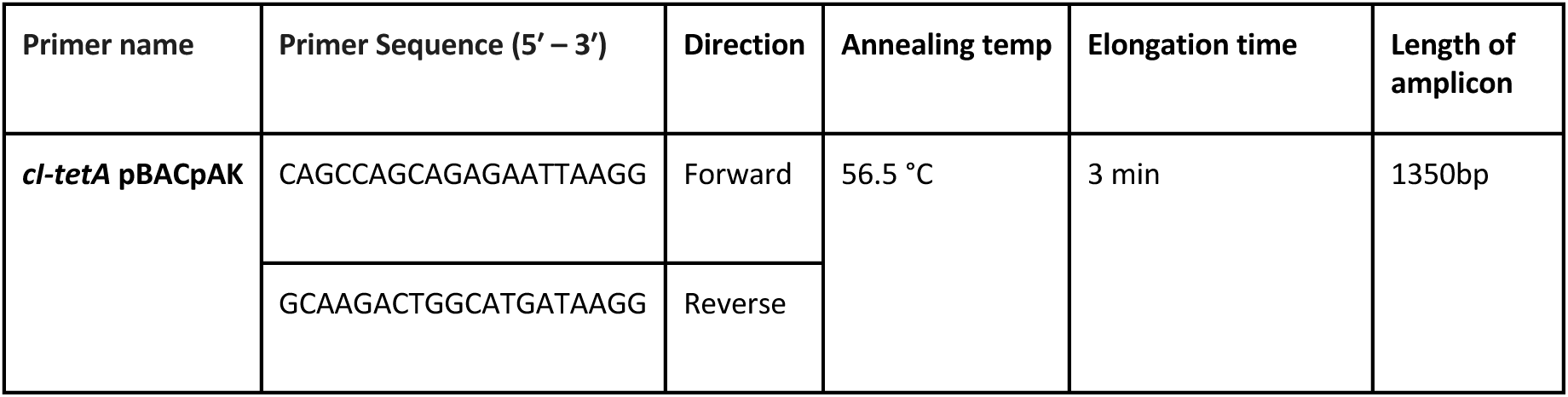
Primers used in this study.

**Supplementary Figure 1:**
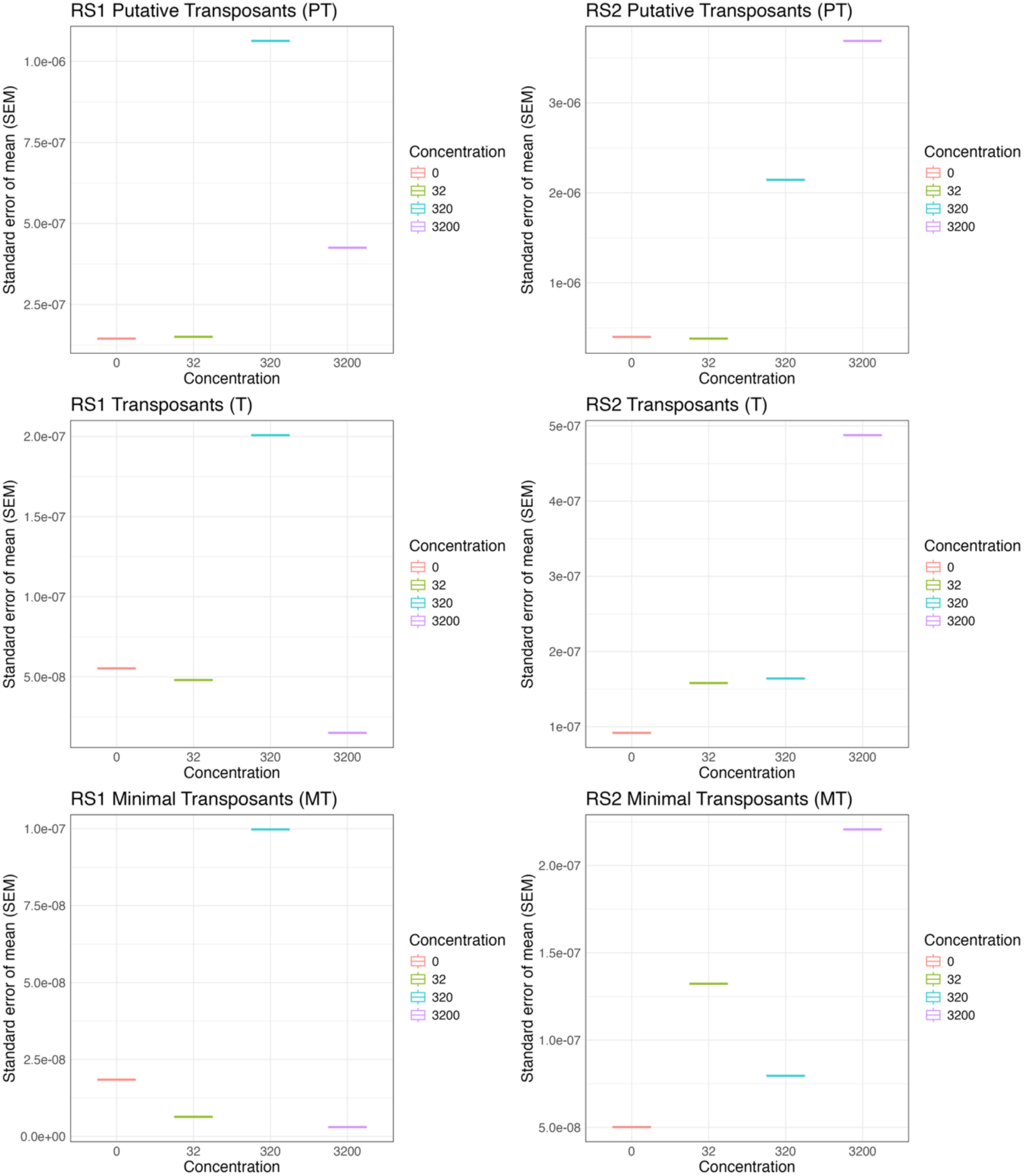
Standard error of the mean (SEM) for putative transposons (PT), transposons (T) and minimal transposons (MT). SEM was calculated using the equation SD/√NR, where SD was the standard deviation and NR was the number of replicates.

**Supplementary Figure 2:**
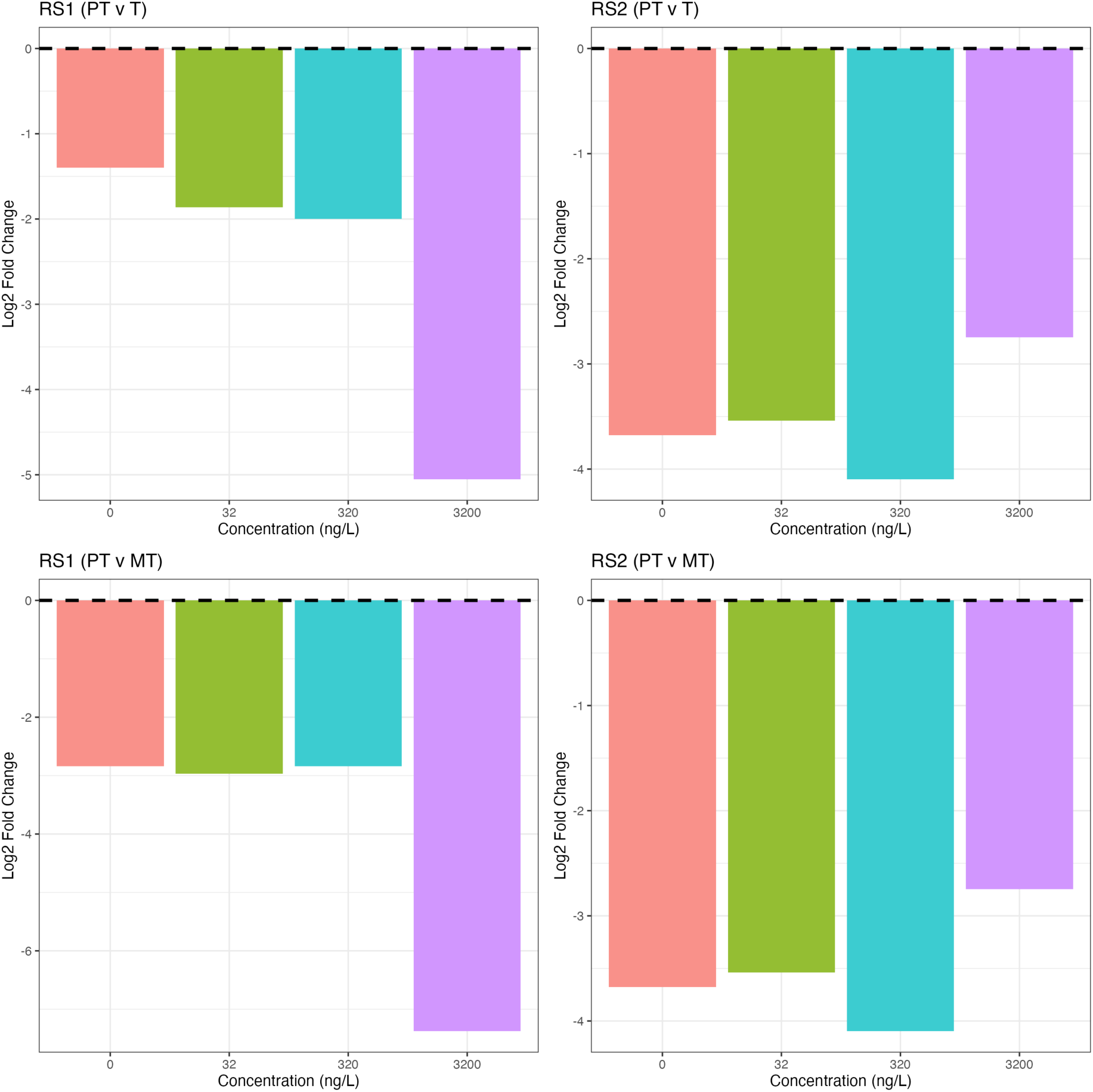
Log_2 f_old change between putative transposons (PT) and transposons (T) and between putative transposons (PT) and minimal transposons (MT) by concentration for RS1 and RS2.

**Supplementary Figure 3:**
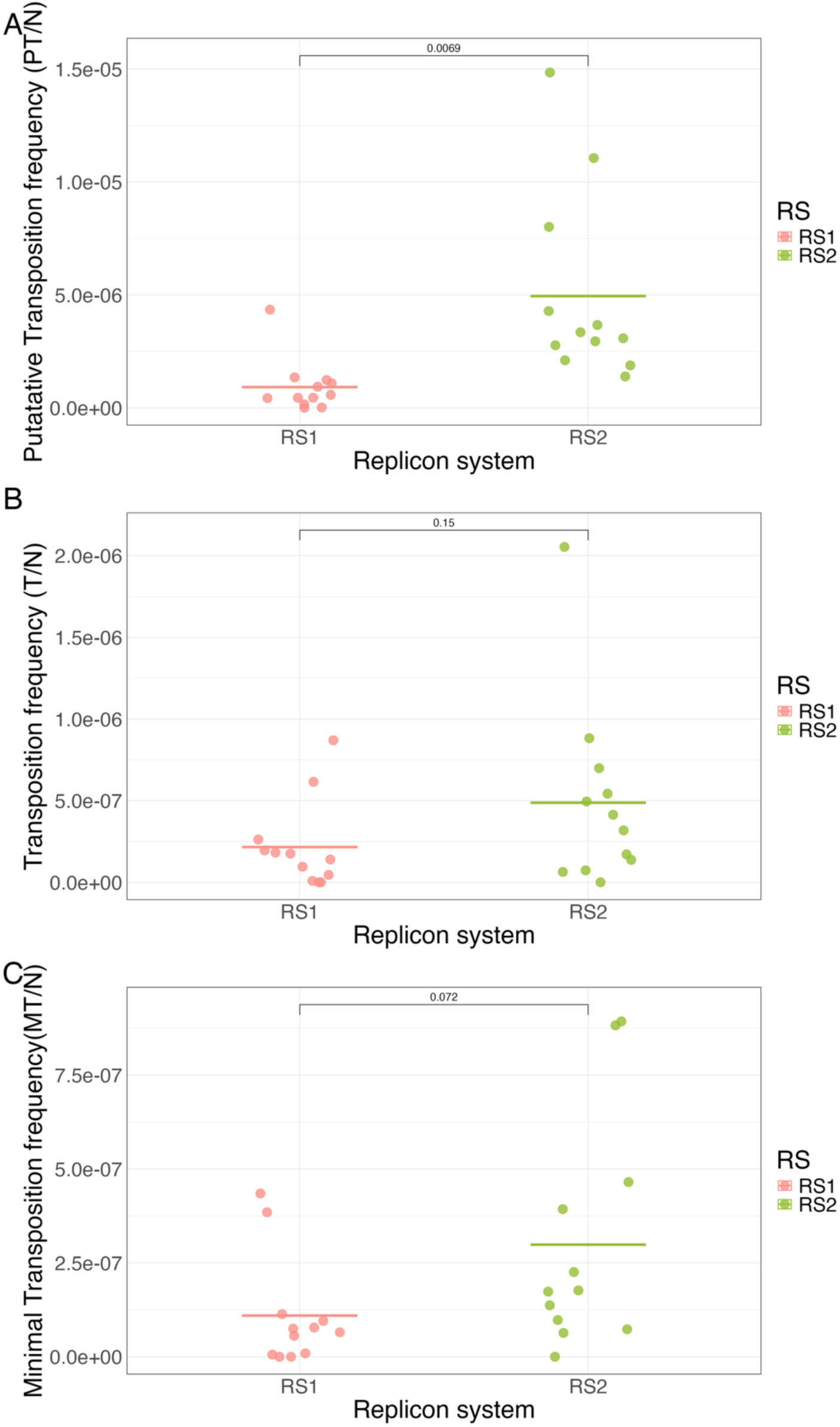
Putative transposition frequency (PT/N) and transposition frequency (T/N) and minimal transposition frequency (MT/N) in replicon system 1 (pBACpAK-COL) vs replicon system 2 (pBACpAK-KAN). Pairwise t-test was run between RS1 and RS2. Horizontal crossbar represent the mean of data points.

**Supplementary Figure 4:**
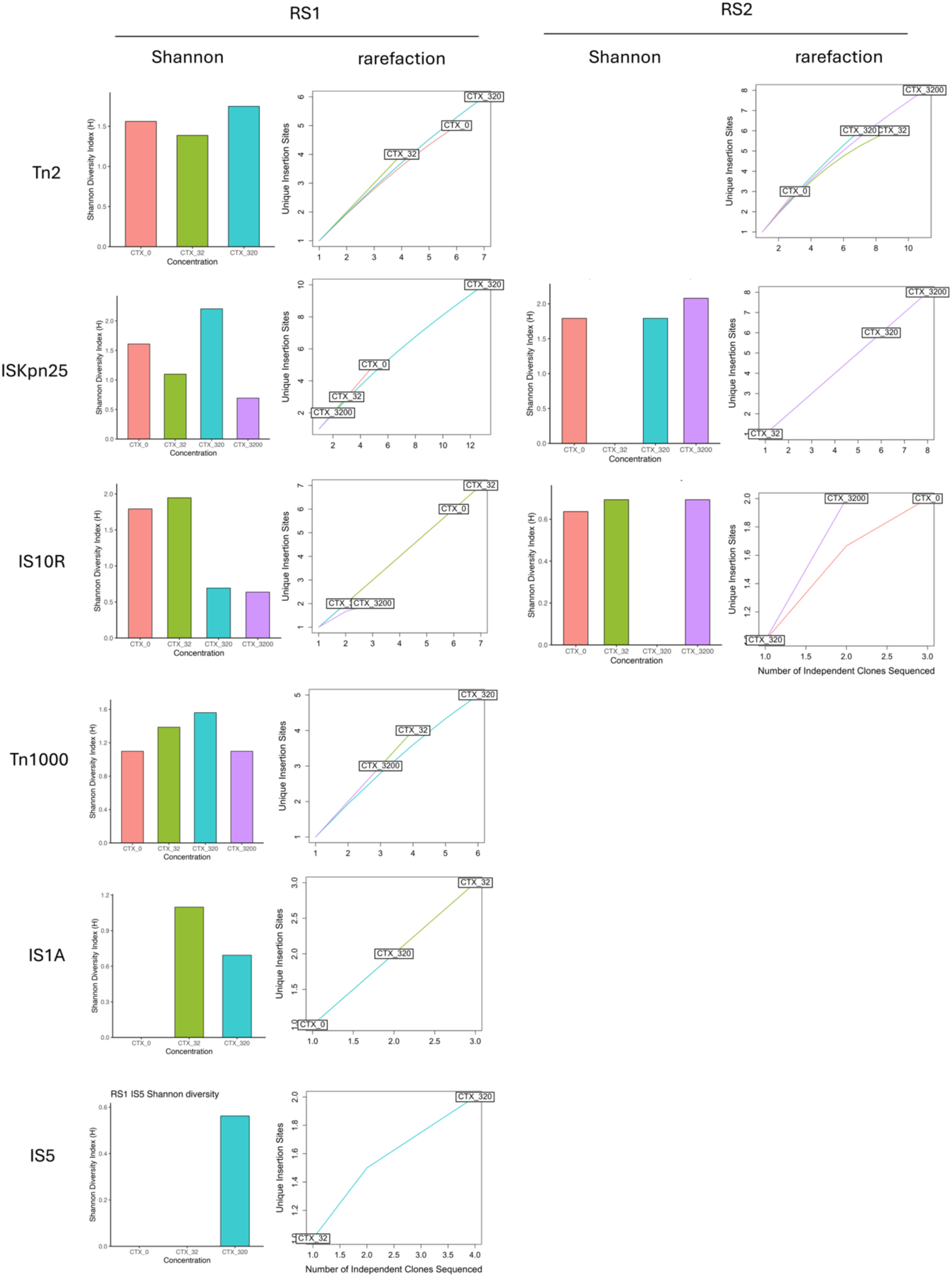
Diversity of insertion site analysis with Shannon diversity and rarefaction.

## References

Ahmad, M. et al. (2025) ‘The role of bacterial metabolism in antimicrobial resistance’, Nature Reviews Microbiology, 23(7), pp. 439–454. doi: 10.1038/s41579-025-01155-0.

Alav, I. et al. (2024) ‘Natural products from food sources can alter the spread of antimicrobial resistance plasmids in Enterobacterales’, Microbiology, 170(8), pp. 1–12. doi: 10.1099/mic.0.001496.

Atsumi, S. and Little, J. W. (2006) ‘Role of the lytic repressor in prophage induction of phage λ as analyzed by a module-replacement approach’, Proceedings of the National Academy of Sciences, 103(12). doi: 10.1073/pnas.0511117103.

Baharoglu, Z. and Mazel, D. (2014) ‘SOS, the formidable strategy of bacteria against aggressions’, FEMS Microbiology Reviews, 38(6), pp. 1126–1145. doi: 10.1111/1574-6976.12077.

Bellanger, X. et al. (2007) ‘Derepression of Excision of Integrative and Potentially Conjugative Elements from Streptococcus thermophilus by DNA Damage Response: Implication of a cI-Related Repressor’, Journal of Bacteriology, 189(4), pp. 1478–1481. doi: 10.1128/JB.01125-06.

Bengtsson-Palme, J. and Larsson, D. G. J. (2016) ‘Concentrations of antibiotics predicted to select for resistant bacteria: Proposed limits for environmental regulation’, Environment International, 86, pp. 140–149. doi: 10.1016/j.envint.2015.10.015.

Boxall, A. B. A. et al. (2012) ‘Pharmaceuticals and personal care products in the environment: What are the big questions?’, Environmental Health Perspectives, pp. 1221–1229. doi: 10.1289/ehp.1104477.

Boxall, A. B. A., Wilkinson, J. L. and Bouzas-Monroy, A. (2022) ‘Medicating nature: Are human-use pharmaceuticals poisoning the environment?’, One Earth. Elsevier Inc., pp. 1080–1084. doi: 10.1016/j.oneear.2022.09.009.

Brouwer, M. S. M. et al. (2020) ‘Mobile colistin resistance gene mcr-1 is detected on an IncI1 plasmid in E. coli from meat’, Journal of Global Antimicrobial Resistance, 23, pp. 145–148. doi: 10.1016/j.jgar.2020.08.018.

Castañeda-Barba, S., Top, E. M. and Stalder, T. (2024) ‘Plasmids, a molecular cornerstone of antimicrobial resistance in the One Health era’, Nature Reviews Microbiology, 22(1), pp. 18–32. doi: 10.1038/s41579-023-00926-x.

Chow, L., Waldron, L. and Gillings, M. R. (2015) ‘Potential impacts of aquatic pollutants: Sub-clinical antibiotic concentrations induce genome changes and promote antibiotic resistance’, Frontiers in Microbiology, 6(AUG), pp. 1–10. doi: 10.3389/fmicb.2015.00803.

Cocker, D. et al. (2025) ‘Environmental hazards from pollution of antibiotics and resistance-driving chemicals in an urban river network from Malawi’, npj Antimicrobials and Resistance, 3(1), p. 85. doi: 10.1038/s44259-025-00149-5.

Darby, E. M. et al. (2022) ‘Molecular mechanisms of antibiotic resistance revisited’, Nature Reviews Microbiology, pp. 1–26. doi: 10.1038/s41579-022-00820-y.

Davies, J., Spiegelman, G. B. and Yim, G. (2006) ‘The world of subinhibitory antibiotic concentrations’, Current Opinion in Microbiology, 9(5), pp. 445–453. doi: 10.1016/j.mib.2006.08.006.

Ding, M. et al. (2022) ‘Subinhibitory antibiotic concentrations promote the horizontal transfer of plasmid-borne resistance genes from Klebsiellae pneumoniae to Escherichia coli’, Frontiers in Microbiology, 13(November), pp. 1–11. doi: 10.3389/fmicb.2022.1017092.

Dwyer, D. J. et al. (2014) ‘Antibiotics induce redox-related physi’, Proceedings of the National Academy of Sciences of the United States of America, 111(20). doi: 10.1073/pnas.1401876111.

Foti, J. J. et al. (2012) ‘Oxidation of the Guanine Nucleotide Pool Underlies Cell Death by Bactericidal Antibiotics’, Science, 336(6079), pp. 315–319. doi: 10.1126/science.1219192.

Galhardo, R. S. et al. (2009) ‘DinB upregulation is the sole role of the SOS response in stress-induced mutagenesis in Escherichia coli’, Genetics, 182(1), pp. 55–68. doi: 10.1534/genetics.109.100735.

Gillings, M. R. (2013) ‘Evolutionary consequences of antibiotic use for the resistome, mobilome, and microbial pangenome’, Frontiers in Microbiology, 4(JAN), pp. 1–10. doi: 10.3389/fmicb.2013.00004.

Goodman, R. N. et al. (2023) ‘Intracellular Transposition of Mobile Genetic Elements Associated with the Colistin Resistance Gene mcr-1’, Microbiology Spectrum. Edited by C. P. Andam, 11(1), pp. 1–13. doi: 10.1128/spectrum.03278-22.

Goodman, R. N., Tansirichaiya, S. and Roberts, A. P. (2023) ‘Development of pBACpAK entrapment vector derivatives to detect intracellular transfer of mobile genetic elements within chloramphenicol resistant bacterial isolates’, Journal of Microbiological Methods, 213(August), p. 106813. doi: 10.1016/j.mimet.2023.106813.

Goswami, C. et al. (2020) ‘Origin, maintenance and spread of antibiotic resistance genes within plasmids and chromosomes of bloodstream isolates of escherichia coli’, Microbial Genomics, 6(4). doi: 10.1099/mgen.0.000353.

Graf, F. E. et al. (2024) ‘Molecular mechanisms of re-emerging chloramphenicol susceptibility in extended-spectrum beta-lactamase-producing Enterobacterales’, Nature Communications, 15(1), p. 9019. doi: 10.1038/s41467-024-53391-2.

Gutierrez, A. et al. (2013) ‘β-lactam antibiotics promote bacterial mutagenesis via an RpoS-mediated reduction in replication fidelity’, Nature Communications, 4. doi: 10.1038/ncomms2607.

Hubbard, A. T. M. et al. (2020) ‘Piperacillin/tazobactam resistance in a clinical isolate of Escherichia coli due to IS26-mediated amplification of bla TEM-1B’, Nature Communications, 11(1), p. 4915. doi: 10.1038/s41467-020-18668-2.

Johansson, M. H. K. et al. (2020) ‘Detection of mobile genetic elements associated with antibiotic resistance in Salmonella enterica using a newly developed web tool: MobileElementFinder’, Journal of Antimicrobial Chemotherapy, pp. 1–9. doi: 10.1093/jac/dkaa390.

Kariuki, S. (2024) ‘Global burden of antimicrobial resistance and forecasts to 2050’, The Lancet, 404(10459), pp. 1172–1173. doi: 10.1016/ S0140-6736(24)01867-1.

Larsson, D. G. J. and Flach, C. F. (2022) ‘Antibiotic resistance in the environment’, Nature Reviews Microbiology, 20(5), pp. 257–269. doi: 10.1038/s41579-021-00649-x.

Lartigue, M. F. et al. (2006) ‘In vitro analysis of ISEcp1B-mediated mobilization of naturally occurring β-lactamase gene blaCTX-M of Kluyvem ascorbata’, Antimicrobial Agents and Chemotherapy, 50(4), pp. 1282–1286. doi: 10.1128/AAC.50.4.1282-1286.2006.

Laureti, L., Matic, I. and Gutierrez, A. (2013) ‘Bacterial responses and genome instability induced by subinhibitory concentrations of antibiotics’, Antibiotics, 2(1), pp. 100–114. doi: 10.3390/antibiotics2010100.

Luria, S. E. and Delbrück, M. (1943) ‘MUTATIONS OF BACTERIA FROM VIRUS SENSITIVITY TO VIRUS RESISTANCE’, Genetics, 28(6), pp. 491–511. doi: 10.1093/genetics/28.6.491.

Maiques, E. et al. (2006) ‘β-lactam antibiotics induce the SOS response and horizontal transfer of virulence factors in Staphylococcus aureus’, Journal of Bacteriology, 188(7), pp. 2726–2729. doi: 10.1128/JB.188.7.2726-2729.2006.

McEwen, S. A. and Collignon, P. J. (2018) ‘Antimicrobial Resistance: a One Health Perspective’, Microbiology Spectrum. Edited by F. M. Aarestrup et al., 6(2). doi: 10.1128/microbiolspec.ARBA-0009-2017.

McGuffie, M. J. and Barrick, J. E. (2021) ‘PLannotate: Engineered plasmid annotation’, Nucleic Acids Research, 49(W1), pp. W516–W522. doi: 10.1093/NAR/GKAB374.

Miller, C. et al. (2004) ‘SOS response induction by β-lactams and bacterial defense against antibiotic lethality’, Science, 305(5690), pp. 1629–1631. doi: 10.1126/science.1101630.

Moubareck, C. A. (2020) ‘Polymyxins and bacterial membranes: A review of antibacterial activity and mechanisms of resistance’, Membranes, 10(8), pp. 1–30. doi: 10.3390/MEMBRANES10080181.

Murray, A. K. et al. (2021) ‘Dawning of a new ERA: Environmental Risk Assessment of antibiotics and their potential to select for antimicrobial resistance’, Water Research, 200, p. 117233. doi: 10.1016/j.watres.2021.117233.

Murray, A. K. et al. (2024) ‘A critical meta-analysis of predicted no effect concentrations for antimicrobial resistance selection in the environment’, Water Research, 266, p. 122310. doi: 10.1016/j.watres.2024.122310.

Murray, C. J. et al. (2022) ‘Global burden of bacterial antimicrobial resistance in 2019: a systematic analysis’, The Lancet. doi: 10.1016/s0140-6736(21)02724-0.

Musicha, P. et al. (2019) ‘Genomic analysis of Klebsiella pneumoniae isolates from Malawi reveals acquisition of multiple ESBL determinants across diverse lineages’, Journal of Antimicrobial Chemotherapy, 74(5), pp. 1223–1232. doi: 10.1093/jac/dkz032.

Nagel, M. et al. (2011) ‘Influence of ciprofloxacin and vancomycin on mutation rate and transposition of IS256 in Staphylococcus aureus’, International Journal of Medical Microbiology, 301(3), pp. 229–236. doi: 10.1016/j.ijmm.2010.08.021.

Negishi, K., Loakes, D. and Schaaper, R. M. (2002) ‘Saturation of DNA mismatch repair and error catastrophe by a base analogue in Escherichia coli’, Genetics, 161(4), pp. 1363–1371. doi: 10.1093/genetics/161.4.1363.

Nicolas, E. et al. (2015) ‘The Tn3-family of Replicative Transposons’, Microbiology Spectrum. Edited by M. Chandler and N. Craig, 3(4). doi: 10.1128/microbiolspec.MDNA3-0060-2014.

Ross, K. et al. (2021) ‘TnCentral: a Prokaryotic Transposable Element Database and Web Portal for Transposon Analysis’, mBio, 12(5). doi: 10.1128/mBio.02060-21.

Sampaio, N. M. V. et al. (2022) ‘Dynamic gene expression and growth underlie cell-to-cell heterogeneity in Escherichia coli stress response’, Proceedings of the National Academy of Sciences of the United States of America, 119(14), pp. 1–12. doi: 10.1073/pnas.2115032119.

Shimoni, Y. et al. (2009) ‘Stochastic analysis of the SOS response in Escherichia coli’, PLoS ONE, 4(5). doi: 10.1371/journal.pone.0005363.

Siguier, P. et al. (2006) ‘ISfinder: the reference centre for bacterial insertion sequences.’, Nucleic acids research, 34(Database issue). doi: 10.1093/NAR/GKJ014.

Singer, A. C. et al. (2016) ‘Review of Antimicrobial Resistance in the Environment and Its Relevance to Environmental Regulators’, Frontiers in Microbiology, 7. doi: 10.3389/fmicb.2016.01728.

Sivertsen, A. et al. (2016) ‘A silenced vanA gene cluster on a transferable plasmid caused an outbreak of vancomycin-variable enterococci’, Antimicrobial Agents and Chemotherapy, 60(7), pp. 4119–4127. doi: 10.1128/AAC.00286-16.

Sołyga, A. and Bartosik, D. (2004) ‘Entrapment vectors--how to capture a functional transposable element.’, Polish journal of microbiology, 53(3), pp. 139–44. Available at: http://www.ncbi.nlm.nih.gov/pubmed/15702911.

Stanton, I. C. et al. (2022) ‘Does Environmental Exposure to Pharmaceutical and Personal Care Product Residues Result in the Selection of Antimicrobial-Resistant Microorganisms, and is this Important in Terms of Human Health Outcomes?’, Environmental Toxicology and Chemistry, 00(00), pp. 1–14. doi: 10.1002/etc.5498.

Stanton, I. C. et al. (2026) ‘Predicting antifungal concentrations that select for resistance: an enhanced approach to establish environmental thresholds’, Environment International, 209, p. 110178. doi: 10.1016/j.envint.2026.110178.

Tansirichaiya, S. et al. (2020) ‘Capture of a novel, antibiotic resistance encoding, mobile genetic element from Escherichia coli using a new entrapment vector’, Journal of Applied Microbiology, pp. 1–11. doi: 10.1111/jam.14837.

Tansirichaiya, S. et al. (2022) ‘Intracellular Transposition and Capture of Mobile Genetic Elements following Intercellular Conjugation of Multidrug Resistance Conjugative Plasmids from Clinical Enterobacteriaceae Isolates’, Microbiology Spectrum. Edited by P. Power, 10(1). doi: 10.1128/spectrum.02140-21.

Tansirichaiya, S. et al. (2025) ‘Derivatization of pBACpAK entrapment vectors for enhanced mobile genetic element transposition detection in multidrug-resistant Escherichia coli’, Access Microbiology, 7(5), pp. 1–11. doi: 10.1099/acmi.0.001013.v3.

Vandecraen, J. et al. (2017) ‘The impact of insertion sequences on bacterial genome plasticity and adaptability’, Critical Reviews in Microbiology, 43(6), pp. 709–730. doi: 10.1080/1040841X.2017.1303661.

Wagner, T. M. et al. (2023) ‘Transiently silent acquired antimicrobial resistance: an emerging challenge in susceptibility testing’, Journal of Antimicrobial Chemotherapy, 78(3), pp. 586–598. doi: 10.1093/jac/dkad024.

World Health Organization (2025) Global antibiotic resistance surveillance report 2025: WHO Global Antimicrobial Resistance and Use Surveillance System (GLASS).

World Health Organization (WHO) (2023) WHO Guidance on waste and wastewater management in pharmaceutical manufacturing with emphasis on antibiotic production. Available at: https://cdn.who.int/media/docs/default-source/wash-documents/burden-of-disease/waste-and-wastewater-management-in-pharma-manufacturing---pub-consult-231214.pdf?sfvrsn=5e5e3acc_3.

Wu, Y., Aandahl, R. Z. and Tanaka, M. M. (2015) ‘Dynamics of bacterial insertion sequences: Can transposition bursts help the elements persist? Theories and models’, BMC Evolutionary Biology, 15(1), pp. 1–12. doi: 10.1186/s12862-015-0560-5.

Xiao, X. et al. (2022) ‘Subinhibitory Concentration of Colistin Promotes the Conjugation Frequencies of Mcr-1-and bla NDM-5 -Positive Plasmids’, Microbiology Spectrum. Edited by T. Polen, 10(2), pp. 1–9. doi: 10.1128/spectrum.02160-21.

Zhou, H. et al. (2024) ‘Environmentally Relevant Concentrations of Tetracycline Promote Horizontal Transfer of Antimicrobial Resistance Genes via Plasmid-Mediated Conjugation’, Foods, 13(11). doi: 10.3390/foods13111787.

